# Genome-scale metabolic models predict diet-and lifestyle-driven shifts of ecological interactions in the gut microbiome

**DOI:** 10.1101/2025.09.23.678088

**Authors:** Georgios Marinos, Karlis Arturs Moors, Malte Rühlemann, Silvio Waschina, Wolfgang Lieb, Andre Franke, Matthias Laudes, Mathieu Groussin, Mathilde Poyet, Christoph Kaleta, A. Samer Kadibalban

## Abstract

Microbiomes and their host environments form complex, interconnected ecosystems. The microbial species within a microbiome, on the one hand, compete for resources, while on the other hand, they exchange vital metabolites to support their survival. These interactions are influenced by the microbial genetic repertoire, environmental conditions, and the availability of nutrients. We developed EcoGS (http://www.github.com/KaletaLab/EcoGS), a metabolic modelling tool designed to predict ecological interactions between pairs of microbes. Applying EcoGS to the microbiomes of two distinct human cohorts revealed a shift from collaborative to exploitative ecological interactions associated with higher dietary intake of simple sugars (glucose and fructose), in diabetic individuals and those living in industrialised lifestyles. On the other hand, the consumption of Cobalamin (vitamin B12), phylloquinone (vitamin K1) and biotin (vitamin B7), among other compounds, was associated with increased collaboration in the gut microbiome. We conclude that the abundance of simple sugars as an energy source reduces the necessity for microbes to cooperate, thereby increasing competition and hostility among microbiome members. Moreover, our study proposes multiple compounds, such as urate, deoxyadenosine, deoxyguanosine, and hypoxanthine, for in vitro validation tests as dietary interventions that have the potential to restore the ecological balance amongst the community. EcoGS serves as a valuable tool for exploring microbiome dynamics and their connections to environmental changes and disease.

## 3. Introduction

The interactions among microbes, as well as between microbes and their host, are complex, yet understanding these relationships is critical for elucidating how the microbiome influences human health. The gut microbiome plays a key role in a variety of diseases, including inflammatory conditions such as Inflammatory Bowel Disease, where current therapies primarily target inflammation rather than directly modulating the microbial community [1]. It also contributes to metabolic disorders, including obesity and diabetes, and influences host immune function and behaviour through the gut-brain axis [2]. Like all microbial communities, the trillions of microbes in the human gut do not act in isolation; they form intricate ecological networks that mediate cooperation, competition, and metabolic exchange. Although each individual’s microbiome is unique, external factors such as lifestyle and environment significantly shape community composition and function [3]. These considerations highlight the importance of investigating microbiome structure and function, as well as the diversity and stability of microbial ecological networks, using advanced computational tools.

Ecological theory considers that microorganisms interact with each other and their environment, resulting in different growth profiles for individual microorganisms in comparison to growth in isolation. Those interactions can be exploitative (amensalism, competition, and antagonism), collaborative (commensalism and mutualism) or neutral [4]. To study those interactions, a basic approach is to model pairs of two species without taking into account the effect of other species. To this end, the co-growth capabilities of two species are directly associated with their ability to compete or collaborate for resources. Specifically, their co-growth rates should be less than their single individual growth rates in case of competition, more in case of collaboration, and the same in case of absence of interaction [5].

To understand interactions within the microbiome, it is important to study the co-occurrence of the involved microbes. Metabolically dissimilar species are more likely to co-occur, as they do not compete for the same resources. Furthermore, metabolic competition is positively associated with phylogenetic relatedness [6, 7]. However, metabolic cooperation can still be predicted. Specifically, cases of close cooperation loops among species, where the products of one species are a nutritional source for other species, have been predicted in communities [5]. Interestingly, cooperation can lead to the stability of the communities in case of nutritional perturbations, while competition ensures stability in case invasive species occur [6].

While the above-mentioned ecological concepts appear theoretically straightforward, inferring them for the gut microbiome is challenging. Importantly, this is because a vast number of microbes cannot be cultivated [8], and the physicochemical environment of the gut that strongly influences microbiome structure is highly variable [9]. Moreover, the human gut microbiome is influenced by human lifestyle and environmental factors, as urbanisation has put pressure on species that were favoured by the pre-agricultural way of life [3]. To this end, a seminal review on the effect of nutritional groups (e.g., amino acids, fats, carbohydrates) and dietary styles (e.g., Western diet) on the abundance of species was published [10]. For instance, the lack of microbiota-accessible carbohydrates in Western-style diets is a reason for species extinction in the gut, which otherwise could produce short-chain fatty acids, leading to detrimental effects, such as dysbiosis, for the host [11].

There is a need for tools that allow categorising ecological interactions within the microbiome as a prerequisite for a targeted manipulation of microbiome structure and function [12]. One key approach that allows for computationally predicting ecological interactions within microbial communities is constraint-based modelling. This methodology builds on genome-scale metabolic reconstructions of the metabolic networks of microbial species that can be derived using automatic tools, like gapseq [13]. Therefore, apart from the genome of the species, nutritional cues, which serve as a proxy for lifestyle and environment, can also be taken into account. Such models can be used for simulating growth profiles of individual microbial species based on linear programming approaches (e.g. flux balance analysis (FBA) [14]). Furthermore, it is possible to simulate complex microbial communities using approaches such as community FBA. This technique merges multiple metabolic models into one, which can then be used in FBA simulations. The MicrobiomeGS2 [15] is such a modelling software, which has been used in reconstructing personalised community microbial models of humans [16] and nematodes [17].

To predict ecological interactions within microbial communities, we present a user-friendly implementation of our approach that predicts ecological interactions between pairs of species within a given community and nutritional input. The software is based on MicrobiomeGS2 and accepts *SBML* metabolic models that are produced by gapseq [13] and loaded with *sybil* [18] as input. It is implemented as an R package available on GitHub (www.github.com/maringos/EcoGS). In our study, we used this tool to predict the ecological interactions of gut microbial species in the microbiomes of a German cohort and to investigate the stability of the proposed interactions by inducing perturbations to the nutritional input of the models. Lastly, we expanded our investigation by comparing and contrasting the prevalence of the ecological relations among individuals from industrialised and non-industrialised settings.

## 4. Methods

### 4.1. Cohort Data

In this study, data from two independent cohorts were analysed. The FoCus, BSP, and SPC cohorts, here referred to collectively as the Focus cohort (PopGen Biobank, Schleswig-Holstein, Germany), included 2,396 individuals and encompassed comprehensive dietary data, 16S rRNA gene sequences of the gut microbiome, as well as clinical data. Metadata variables included age, gender, body mass index (BMI), smoking status, diabetes, inflammatory bowel disease (IBD), and other relevant health parameters. The cohort consisted of 1,329 females and 1,067 males, with 274 individuals having diabetes (Supplementary figure S1). The data of the 108 individuals diagnosed with IBD were filtered out before the downstream analysis, as IBD has a substantial influence on the gut microbiome structure and dynamics [19]. The 16S rRNA sequences of the gut microbiome of the FoCus cohort were mapped against a reference database of 822 human gut microbial genomes derived from the AGORA resource [20] using USEARCH version v10 [21]. This mapping identified 409 microbial strains with varying abundances across the cohort. The second cohort in this study comprised 1,229 participants as a part of the Global Microbiome Conservancy (GMbC) [22]. This cohort consisted of 789 individuals from non-industrialised communities and 440 from industrialised communities (Supplementary figure S2). For further analysis, we used both the metagenomic data generated as well as whole-genome assemblies. The metadata included age, gender, and body mass index (BMI).

### 4.2. Creating and simulating the community models

gapseq (version 1.2) [13] was used to create genome-scale metabolic models for both the reference (AGORA) microbial genomes and the metagenomic bins of the GMbC cohort, utilising a standardised North German diet [16]. Thereafter, the biomass reaction of the metabolic models was optimised based on flux balance analysis using the R [23] package *sybil* [18], and the achieved baseline growth rate was recorded. Subsequently, the models were joined in pairs using the R package MicrobiomeGS2. After joining each pair of models, and to simulate the community growth, the objective coefficients of each sub-model were set to one; the objective coefficients of other reactions were set to 0. In the next step, following a published approach, we set the following coupling constraints: c = 400 mmol.g ^-1^.hr^-1^ as proposed originally [24] and a more relaxed coupling u = 1e-6 mmol.g ^-1^.hr^-1^ Then we performed flux balance analysis, as originally designed in MicrobiomeGS2. We explicitly recorded the exchange reactions whose fluxes predict the exchange of compounds among the sub-models when one flux is positive and the other is negative, respectively. To ensure the quality of the results, we recorded for each case if both submodels could grow (cut-off 1e^-6^) and if each optimisation process was successful. Both the simulations of the stand-alone models and of the joined models were conducted using the academic version of the linear solver IBM ILOG CPLEX through its R interface, cplexAPI [25], and using the Sybil method hybbaropt. Both simulations run in parallel using multi-core computing based on PSOCK clusters in R.

### 4.3. Inference of the ecological interactions

For each of the 409 microbial species found in participants in the FoCus cohort and the 483 vspecies found in participants of the GMbC cohort, we compared the predicted alone-growth rate (AG) with its co-growth rate (CG) when joined with each of the other microbial species belonging to the same cohort. A change in co-growth rate was considered significant if it differed from the alone-growth rate by at least 1% 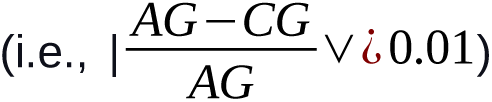 (Figure 1 and Table 1). Based on these comparisons, we constructed a symmetrical matrix representing the inferred ecological interactions between species pairs for each of the two cohorts, encompassing a total of 83,436 pairs for the FoCus microbes (Supplementary Table S1) and 116,403 pairs for the GMbC microbes (Supplementary Table S1). The number of pairs can be calculated as follows

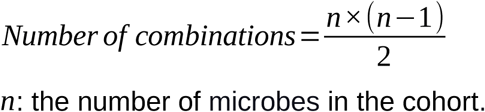

**Figure 1:**
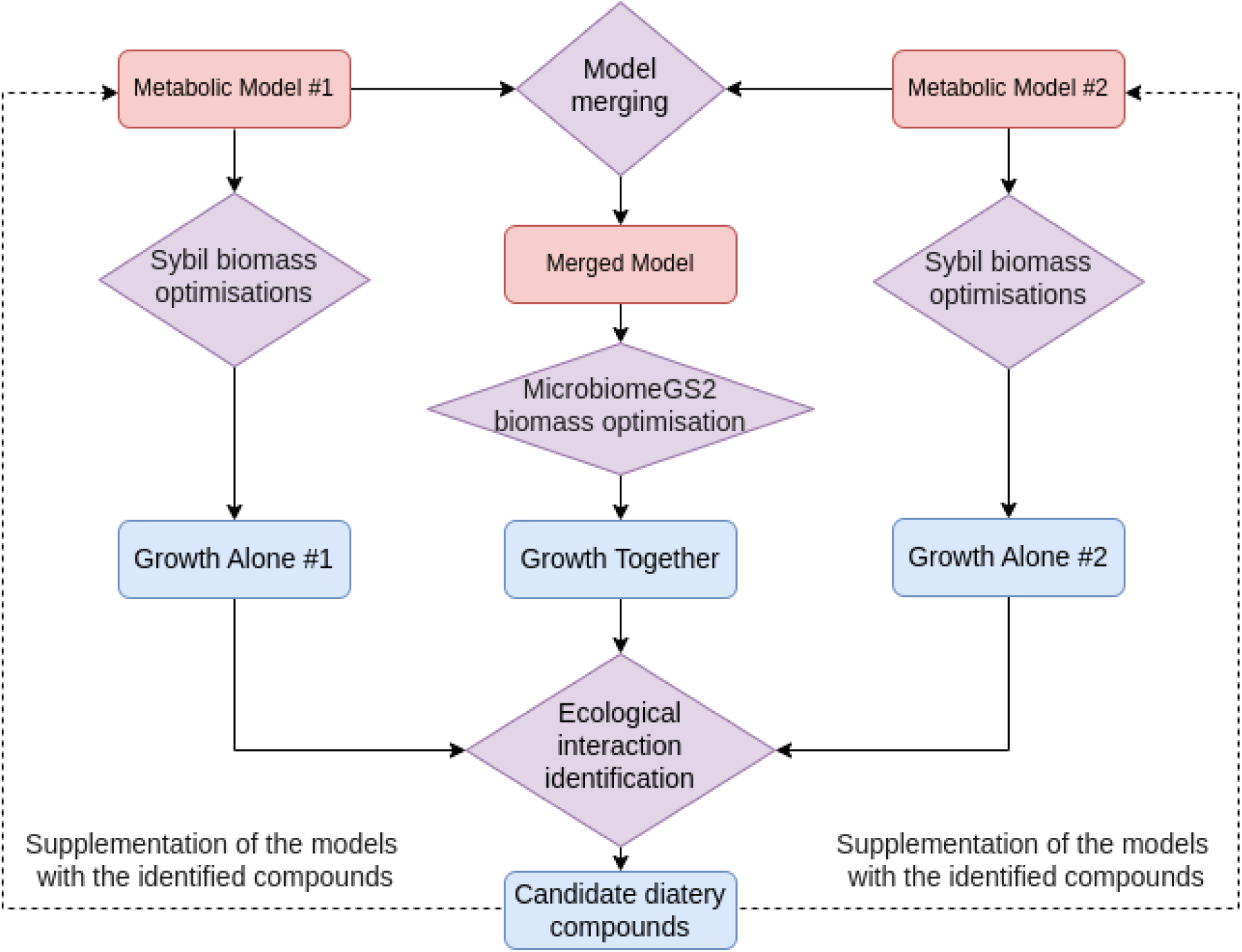
Diagram representation of the EcoGS pipeline: Red boxes represent metabolic model data, blue boxes represent predictions extracted from the models, purple diamond shapes represent operations and functions included in the package, continuous arrows represent the data flow between each step, and dashed arrows represent the feedback of candidate intervention data for predicting their influence.

**Table 1:** Definition of the ecological interactions for two species, a and b.

|  |  |  |  |  |  |
| --- | --- | --- | --- | --- | --- |
| Model #a growth comparison | Model #a interaction effect | Model #b growth comparison | Model #b interaction effect | Ecological interaction | Interaction Category |
| $CG_a > AG_a + m$ | benefit | $CG_b > AG_b + m$ | benefit | Mutualism | Collaborative |
| $AG_a - m \leq CG_a$<br>$\geq AG_a + m$ | no change | $CG_b > AG_b + m$ | benefit | Commensalism | Collaborative |
| $AG_a - m \leq CG_a$<br>$\geq AG_a + m$ | no change | $AG_b - m \leq CG_b \geq AG_b + m$ | no change | Neutralism | Neutral |
| $CG_a > AG_a + m$ | benefit | $CG_b < AG_b - m$ | harm | Antagonism | Exploitative |
| $AG_a - m \leq CG_a$<br>$\geq AG_a + m$ | no change | $CG_b < AG_b - m$ | harm | Amensalism | Exploitative |
| $CG_a < AG_a - m$ | harm | $CG_b < AG_b - m$ | harm | Competition | Exploitative |
| CG: co-growth, AG: Alone growth, m: margin = (Alone growth)/100 |  |  |  |  |  |

Next, we categorised each microbial species pair into one of the possible six ecological interactions (Table 1). Thereafter, we counted the number of pairs belonging to each ecological interaction in each microbial community (4,236 communities in the two cohorts combined) and weighed those counts by the abundance of the species in each pair.

Let the abundances of species *S1, S2, S3, S4, S5*, etc. be denoted as *A1, A2, A3, A4, A5*, etc. For a given ecological interaction type *E*, the total number of species pairs exhibiting this interaction in the community is NE, represented as: (S1, S2), (S1, S3), (S3, S5), etc. The weighted estimate of this ecological interaction, W_NE_, can be calculated in two ways:

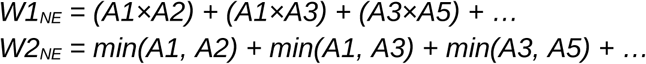

W1_NE_ takes into account the probability of two species encountering each other in the community as the multiplication of their abundances. W2_NE_ takes into account that the limiting factor in the metabolic exchanges between the two species is the minimum abundance of the two. EcoGS allows the user to choose the weighing method. Here, we opted to use the second approach (W2_NE_), as it showed stronger signals in the results.

To address the inherent codependency in compositional data, a logarithmic transformation of the ratio between pairs of variables is often applied [26]. In this study, the proportions of different ecological interactions are compositional data. In the context of the six microbial ecological interactions, this approach yielded 15 log-transformed ecological interaction ratios (EIRs), such as *log(Antagonism/Competition)*, *log(Commensalism/Mutualism)*, and so forth.

The log-transformed proportional data of the ecological interaction ratios per microbial community (with pseudoanonymised IDs) are in supplementary table S1 for the FoCus cohort and in supplementary table S1 for the GmBC cohort.

### 4.4 Phylogenetic inference of the GMbC species

Phylogenetic trees were reconstructed using SGB representative genomes with the identify, align, and infer functions of GTDB-Tk v2.3.2 and GTDB Release R07-RS207 [27, 28]. The bacterial and archaeal trees were rooted with representatives of the Patescibacteria and Nanoarchaeota, respectively, and joined at the root to yield a combined phylogenetic tree encompassing both domains.

### 4.5 Dietary interventions

To predict the effects of dietary interventions on ecological interactions within the gut microbiome of the FoCus cohort, simulations were performed for the supplementation of 105 individual nutrients (metabolites). Each simulation involved modifying the microbial diet in *gapseq* models by adjusting the lower bounds of the exchange reaction for the respective metabolite. The supplementation amount for each nutrient was set to two times the value of the standardised North German diet (Supplementary Table S1). For each intervention, the ecological interaction prediction pipeline was applied to microbial pairs independently, generating 105 ecological interaction matrices with the same dimensions and structure as the original pre-intervention matrix. Thereafter, post-intervention ecological interaction ratios per participant were calculated.

### 4.6 Statistical analysis

For the FoCus cohort, linear models (LMs) were employed to assess associations between the ratios of ecological interactions (EIRs) and four key clinical variables: age, body mass index (BMI), gender, and smoking behaviour (measured as cigarettes per day), as well as with diabetes. The model structure was specified as:

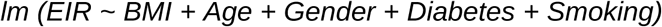

Additionally, EIRs were tested for associations with individual dietary metabolites, which were inferred from dietary questionnaires. Each metabolite was analysed separately, adjusting for the same four clinical covariates:

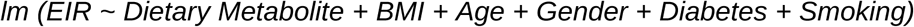

Similarly, for the GMbC cohort, linear models were applied to examine the associations between EIRs and age, BMI, gender, and lifestyle (classified as industrialised or non-industrialised):

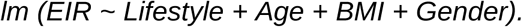

Following the metabolic supplementation simulations, the effect of each supplementation on the EIRs was evaluated by comparing the EIRs before and after supplementation using a paired Wilcoxon signed-rank test. Finally, the phylogenetic distances between each pair of microbes in the GMbC cohort were compared across the six ecological interaction categories using pairwise Wilcoxon signed-rank tests. False discovery rate (FDR) [29] correction was applied to the resulting *P*-values to account for multiple testing after each statistical test. Adjusted *P*-values less than α = 0.05 were considered statistically significant.

### 4.7 Software

The community model simulations were conducted in a high-performance computing environment in an R environment (version 4.4.1) [23]. Specifically, the following R packages were used: MicrobiomeGS2 [15] version 0.2.0, cplexAPI [25] version 1.4.0 (along with the IBM ILOG CPLEX Optimisation Studio / Academic Initiative version 22.10), *sybil* [18] version 2.2.1, doParallel [30] version 1.0.17, stringr version 1.5.1, [31] lattice version 0.22-6, Matrix version 1.7-1, iterators version 1.0.1 4, data.table version 1.17.8, and foreach[32] version 1.5.2.

All statistical analyses were conducted with R (version 4.5.0) within the RStudio environment [33]. Plots were created with the ggplot2 R package (version 3.5.2) [34] and R base plotting functions. The results of all statistical tests were determined to be significant below a threshold of α=0.05. P-values were adjusted for multiple testing using FDR [29].

## 5. Results

### 5.1. Ecological interaction and metabolic exchange among microbial communities and across phylogeny

While ecological interaction categories (amensalism, competition, antagonism, commensalism, mutualism, and neutralism) address the relationship between the interacting species, it is also noteworthy to examine the interaction effect on each model (microbial species) independently (harmful, beneficial, and neutral). The ecological interactions among the gut microbial communities of both cohorts have considerably higher proportions and higher variances of exploitative ecological interactions in comparison to the collaborative and neutral interactions (Figures 2. A and 2B). Furthermore, species pairs with exploitative interactions tend to have higher phylogenetic distances with a bimodality, suggesting that very closely related species also tend to have exploitative interactions, while collaborative and neutral interactions tend to have lower to intermediate phylogenetic distances as observed in the GMbC cohort (Figure 2C). In order to evaluate the differences in the ecological interactions between different microbial genera, the 18 most abundant genera in the gut communities of the FoCus cohort participants were selected.

**Figure 2:**
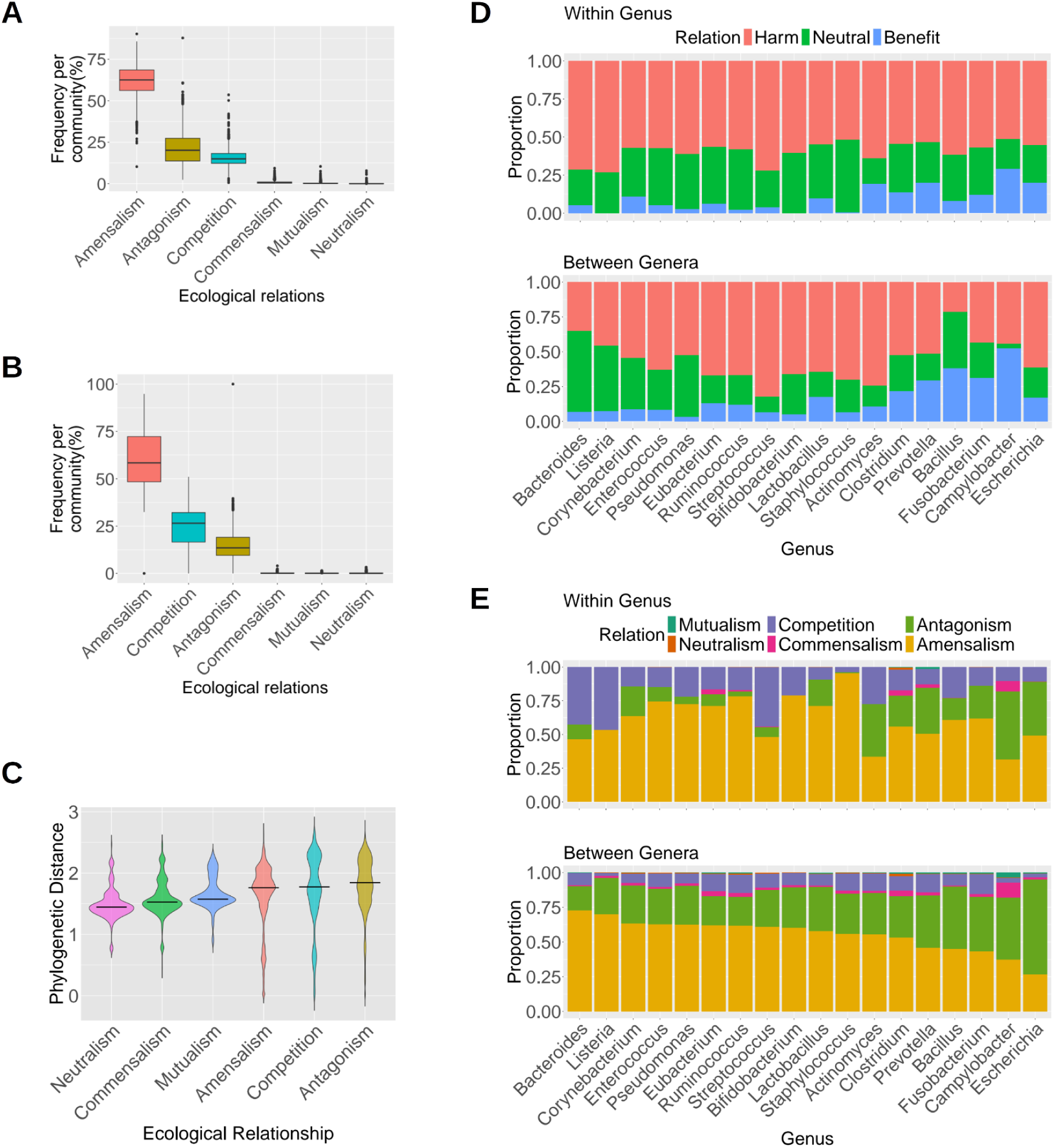
Ecological interactions among microbial communities. A. The frequencies of the six types of ecological interactions among the gut microbiome of the FoCus cohort; the x-axis is ordered by the median of the ecological interaction frequencies across all communities. B. Similar to A. for the GmbC cohort microbial communities, the colours correspond to those in A. C. A violin plot representing the distribution of the phylogenetic distances between pairs of microbes among the six different types of ecological interactions for microbes found in the GMbC cohort participants. The black line in the middle of each box represents the median value. The violins are sorted by median on the x-axis. The colours correspond to those in A. and B. D. Bar plots depicting the interaction outcome for the 18 most abundant genera in the FoCus cohort. The top part shows the outcome of interacting with species from different genera. The bottom part shows the outcome of interacting with species belonging to the same genus. The genera are ordered on the X-axis to correspond to the order in E. E. Bar plots depicting the ecological interaction proportions for the 18 most abundant genera in the FoCus cohort. The top part shows the ecological interactions between microbes of the corresponding genus and different genera. The bottom part shows the ecological interactions between species belonging to the same genus. The genera are ordered on the X-axis in correspondence to their frequency of amensalistic interaction with other genera.

The interaction effect of each model was estimated within each genus and between genera. The former takes into account the interactions among microbial species belonging to each of the genera, and the latter between the species of each of the genera and the species of all other genera. (Figure 2D). Similarly, the proportion of the ecological interactions (considering the interaction effect on both microbes in the pair) was estimated for microbial pairs within each genus and between genera (Figure 2E).

The proportions of interaction effects, on each model, vary between the different genera; the interactions among species from the same genus produce more neutral and less beneficial effects than interactions with species from different genera. The ecological interaction categories also vary between different genera. Higher antagonism, lower competition and higher commensalism are observed in interactions between species belonging to different genera in comparison to interactions within the same genus.

Not only does *Escherichia* have the highest frequency of antagonistic interactions in comparison to other genera, but also, antagonism comprises the highest proportion of all types of ecological interactions between *Escherichia* and other genera. Moreover, those antagonistic interactions are mostly harmful to *Escherichia* (Figures 2D and 2E and supplementary table S2). *Campylobacter, Clostridium*, and *Eubacterium,* on the other hand, have the highest proportions of collaborative interactions with other genera and within the same genus. Whereas *Fusobacterium*, *Prevotella*, *Ruminococcus*, and *Eubacterium* have relatively higher competition against other genera (Figure 2E and Supplementary Table S2). On the other hand, *Bacteroides* species showed a highly amensalistic trend between genera and a competitive trend within the genus. *Staphylococcus* exhibits a very high proportion of amensalistic interactions within the same genus.

Furthermore, we observed a positive association between the number of predicted metabolic exchanges among microbial pairs in the GMbC cohort and their phylogenetic distance; this relationship remained significant even when considering only amino acid exchanges (Figures 3A and 3B). In contrast, a similar pattern was not evident in the FoCus cohort. When comparing exchange frequencies within the same genus versus across different genera—after normalising by the number of interacting pairs in each category—we found no strong differences for either total metabolic exchanges or amino acid-specific exchanges. The only notable exception was *Listeria* and *Prevotella*, which exhibited substantially more amino acid exchanges within their respective genera than with other genera.

**Figure 3:**
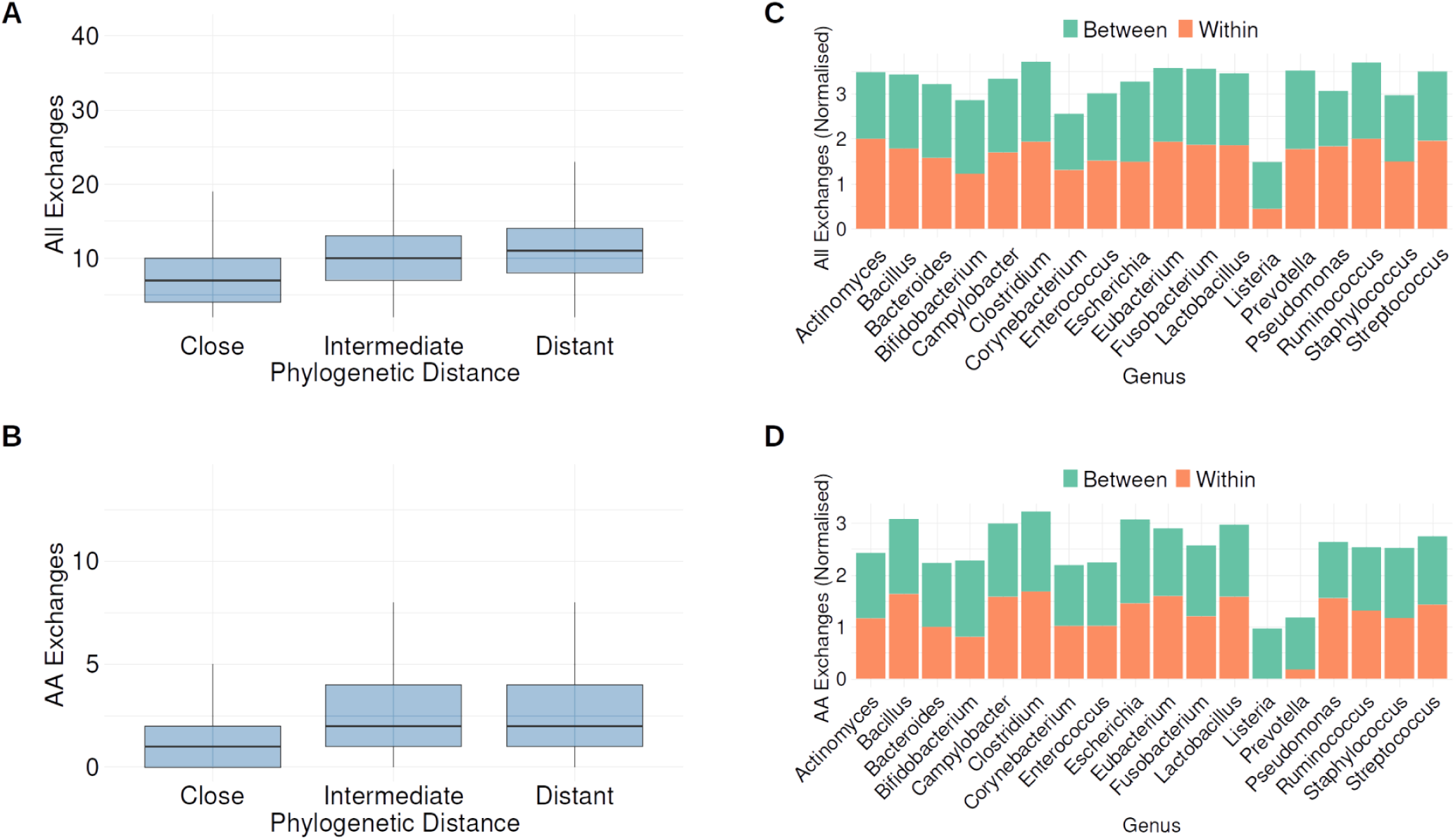
The number of metabolic exchanges between microbial pairs across phylogeny and genera. A. The number of predicted exchanged metabolites between microbial pairs on the Y-axis, across three binned categories of phylogenetic distance between the microbes in each pair on the X-axis. B. Same as A., but only counting the number of amino acid exchanges between the microbes in the pairs. C. A bar plot of the normalised rates of predicted metabolic exchanges between microbes in pairs within the same genus in orange and those belonging to different genera in green. D. Same as C., but only counting the normalised rates of amino acid exchanges between the microbes in the pairs.

### 5.2. Associations between phenotypic and dietary data and the ecological interaction ratios

The linear regression models revealed significant associations between ecological interaction ratios (EIRs) and phenotypical data from both cohorts (e.g. a positive association between diabetes and the proportion antagonism/mutualism and antagonism/commensalism, Figure 4A). Such a positive estimate reflects an increase in the numerator (antagonism) relative to the denominators (mutualism and commensalism).

**Figure 4:**
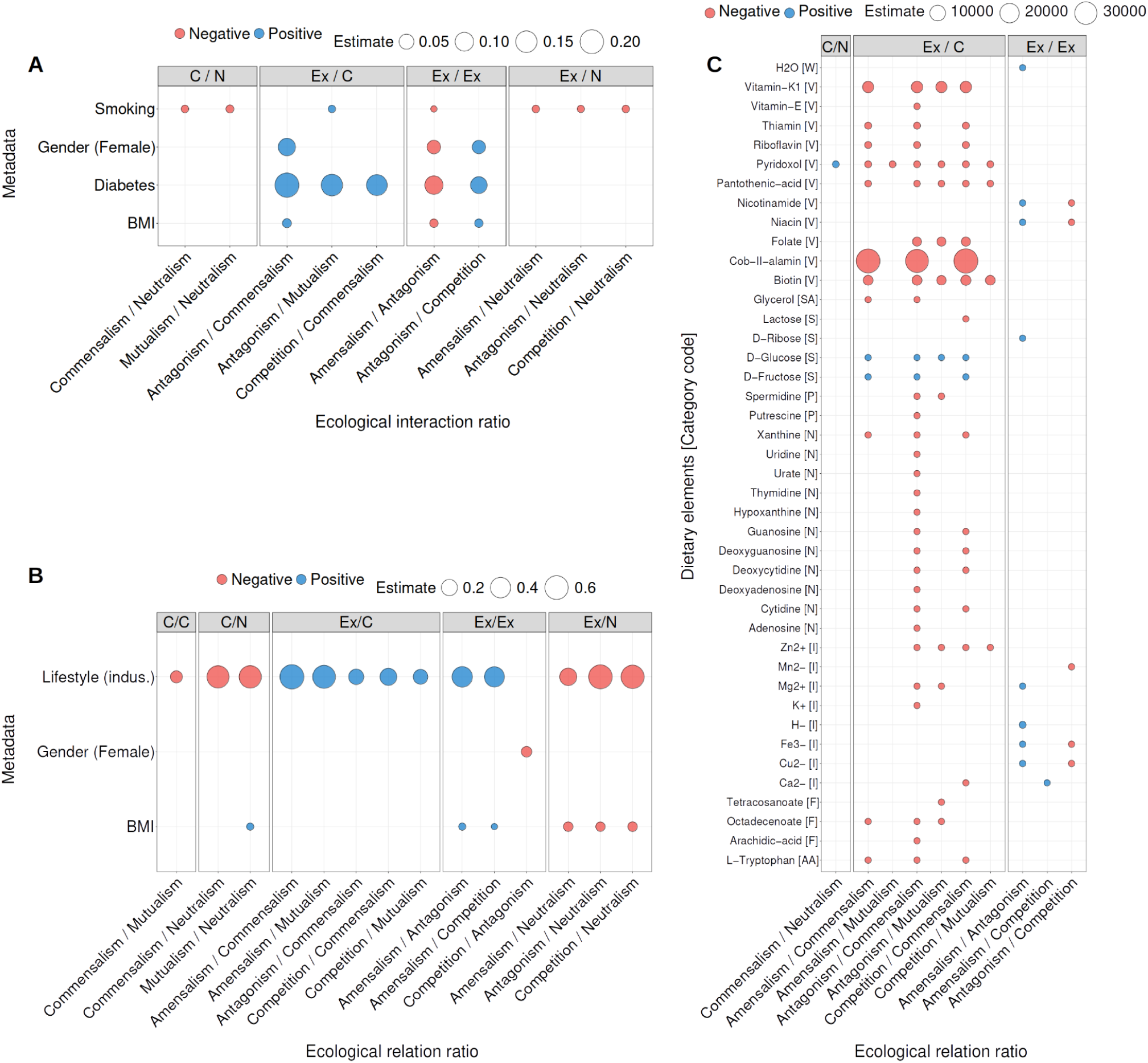
Associations between phenotypic and dietary data with ecological relation ratios. A. Associations between metadata of the FoCus cohort participants and the ecological interaction ratios in their gut microbiomes. Each dot represents one significant association resulting from (LM) between four phenotypic (metadata) on the Y-axis and the ecological interaction ratios on the X-axis (log transformed). The colour represents the directionality of the association, blue for positive and red for negative associations. The size of the dots depicts the LM estimate (coefficient of correlation). The interaction ratios are divided into three panels to distinguish between three types of interaction ratios: Exploitative-exploitative, Exploitative-collaborative, and Collaborative-collaborative. To enhance readability, we grouped the compounds into categories (W for water, V for vitamin, SA for sugar alcohol, S for sugar, P for polyamines, N for Nitrogen-containing compounds, I for ions, F for fatty acids, and AA for amino acids). B. Associations between dietary elements from the diet of the FoCus cohort (y-axis) and the estimated ecological relation ratios of their gut microbiomes (x-axis). The colour and size of the dots and the two panels are as in A (there are only two panels because there are no significant associations with the collaborative-collaborative type of interaction ratios). C. Associations between metadata of the GMbC cohort participants (x-axis) and the ecological interaction ratios in their gut microbiomes (y-axis). The colour and size of the dots and the two panels are as in A.

Similarly, a negative estimate reflects a decrease in the numerator relative to the denominator (e.g. lifestyle vs. antagonism/neutralism, Figure 4B).

In the FoCus cohort, antagonism proportion relative to commensalism and mutualism is significantly increased in the gut microbiome of female participants in comparison to males (P values in supplementary table S2). However, we don’t observe the same effect in the GMbC cohort (Figures 4A and 3B). Smoking, on the other hand, was associated with higher neutral interactions relative to collaborative and exploitative ones and higher antagonism relative to mutualism (Figure 4A). A positive association between BMI and the proportion of antagonism relative to commensalism was observed in the FoCus cohort. However, in the GMbC cohort, we only observe a decrease in exploitative interactions relative to neutralism in association with BMI (Figures 4A and 4B). Moreover, not only is there a relative increase in antagonism relative to collaborative interactions associated with diabetes, BMI, smoking and in females, but antagonism is also increased relative to the two other exploitative interactions (amensalism and competition) in association with the same four phenotypical data types.

Furthermore, we observed an increase in amensalism, competition and antagonism at the expense of collaborative ecological interactions in individuals living in industrialised communities in comparison to individuals living non-industrialised lifestyles in the GMbC cohort. However, the proportions of exploitative and collaborative interactions relative to neutral ones are significantly lower in industrialised populations. (Figure 4B)

Furthermore, we tested for associations between EIRs and different nutrients from the diet of the FoCus cohort. 32 of the tested dietary elements showed a negative association with at least one exploitative relative to collaborative ecological interaction types, including vitamins, ions, fatty acids, nucleotides, polyamines, as well as the amino acid Tryptophan. The majority of those dietary elements had significant associations with increased commensalism. The strongest associations were observed between Cobalamin (vitamin B12), phylloquinone (vitamin K1) and biotin (vitamin B7) with reduced exploitative relative to collaborative interaction ratios. On the other hand, we observed a significant increase in all three exploitative interactions relative to commensalism associated with higher dietary glucose and fructose intake, as well as an increase in antagonism relative to mutualism in association with glucose intake (Figure 4C).

### 5.3. Dietary interventions to alter the ecological interaction distributions along gut microbiomes

To identify dietary components that could alter the ecological interaction ratios and restore the observed shifts associated with diseases such as diabetes, we assessed the impact of individual nutrients on predicted interaction ratios (see Methods). Among the tested metabolites, 19 showed a significant effect on these ratios. Supplementation with certain compounds, including the fatty acid octadecanoate, the nucleotide cytidine, seven purines (deoxyadenosine, guanosine, xanthine, urate, deoxyguanosine, hypoxanthine, and adenosine), and two vitamins (thiamin B1 and riboflavin B2), was associated with an increase in collaborative interactions (mutualism and commensalism) relative to exploitative interactions (antagonism and competition).

Notably, supplementation with D-glucose and D-fructose increased collaborative interactions, whereas L-tryptophan supplementation increased exploitative interactions (Figures 4C and 5; see Discussion).

**Figure 5:**
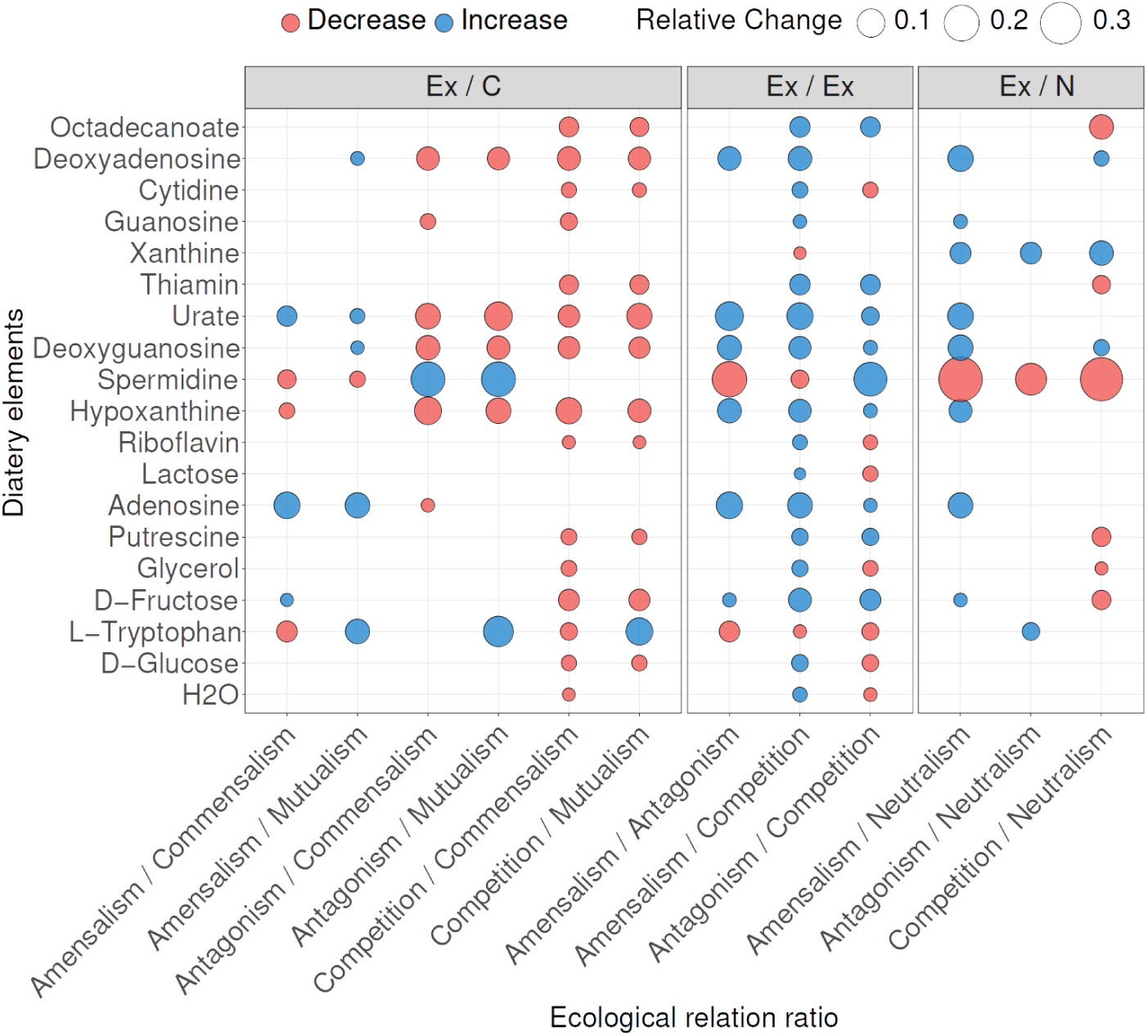
The effect of dietary metabolic intervention (Y-axis) on the ecological relation ratios (X-axis), the dots represent the significant difference between the interaction ratios with and without the intervention, detected with the Wilcoxon signed-rank test. The colour of the dots represents the direction of the change caused by the intervention, red for a decrease in the interaction ratio and blue for an increase. The size of the dots shows the fold change caused by an intervention, depicted as the median of all relative changes among all communities. The two panels discriminate between exploitative-collaborative and exploitative-exploitative interaction ratios.

Within the exploitative category, several interventions altered internal ratios, with most increasing amensalism relative to antagonism and competition, and in some cases, increasing antagonism relative to competition. Notably, L-tryptophan produced an opposite pattern. Additional shifts were observed between exploitative and neutral interactions. Some metabolites promoted amensalism over neutralism, some promoted neutralism over competition, and only spermidine increased neutralism relative to all three exploitative interaction types (Figure 5).

## 6. Discussion

In this study, we introduce *EcoGS*, an open-source R-based computational tool designed to infer ecological interactions among microbial species from genome-scale metabolic models. By simulating the growth of each species both alone and together with another, *EcoGS* predicts interaction types: Exploitative interactions (amensalism, antagonism, and competition), collaborative interactions (commensalism and mutualism) and neutral interactions and summarises them at the community level. This framework offers a scalable and user-friendly approach for exploring microbial ecology in complex communities and is freely available as an R package (www.github.com/KaletaLab/EcoGS).

Applying *EcoGS* to gut microbiomes from two distinct human cohorts, the FoCus cohort from Germany and the GmbC cohort encompassing both industrialised and non-industrialised populations, yielded key insights into how host-related metabolic factors, lifestyle, and diet shape microbial interaction networks. Notably, in both cohorts, we observe higher proportions of exploitative interactions than collaborative ones (Figures 2A and 2B). This pattern is in line with the previously observed frequent co-occurrence among competing species, suggesting that shared habitat filtering drives microbiome structure [7]. It also aligns with controlled experiments showing that competitive and amensal interactions dominate in synthetic human gut communities (comprising ∼68 % of all pairwise interactions, vs. ∼5 % cooperative ones) [12], as well as with recent data showing predominance of negative interactions in natural gut isolates [35].

The phylogenetic distance of interacting species revealed more exploitative interactions between distant species (Figure 2C). However, we also observe a bimodal distribution of pairwise distances among microbes engaged in exploitative interactions, suggesting that those interactions also occur among closely related and distantly related taxa (Figure 2C). In addition, a positive correlation between the number of metabolic exchanges and phylogenetic distance (Figures 3A and 3B) supports the idea that closer species are metabolically similar, share similar enzyme systems, carbon source preferences, and oxygen requirements. Due to this niche overlap and evolutionary constraints on metabolic complementation, they tend to engage in less frequent metabolite sharing [36].

In the FoCus cohort, we observe an increase in antagonism relative to commensalism in individuals with a higher body mass index (BMI) and in individuals with diabetes, as well as an increase in antagonism relative to mutualism in diabetic individuals. This shift away from collaborative or neutral interactions may reflect a disrupted or less cooperative microbiome in metabolically compromised states, potentially reinforcing disease-associated dysbiosis. However, in the GMbC cohort, there is only an increase in neutralism over other types of interactions in association with BMI. This could be due to the structure of the cohort, which includes individuals from different lifestyles. With this, our study extends findings from a previous study [37] across independent populations. In that previous study, it was found that elevated blood glucose levels and the presence of diabetes were associated with a marked increase in exploitative microbial interactions, particularly antagonism.

Moreover, in the GMbC cohort, microbiomes from participants with industrialised lifestyles displayed a dual trend: A rise in exploitative interactions over collaborative ones, alongside an increase in neutral interactions. This suggests that industrialisation not only tilts microbial communities toward more competitive dynamics but also introduces ecological stasis or tolerance, where species coexist without significant interaction, a possible reflection of ecological simplification driven by uniform diets, reduced microbial diversity, or environmental exposures associated with industrialisation [38]. Together, these findings indicate that both metabolic disease and industrialisation converge on a common ecological footprint: reduced cooperation and increased exploitation within the gut microbial community. Both diabetes and industrialisation often imply higher availability of simple carbohydrates, highlighting a potential mechanism by which modern diets contribute to microbiome instability and drive metabolic dysregulation, as the availability of simple, rapidly metabolised energy sources reduces the need for microbial cooperation.

In the human gut, it is suggested that species have distinct roles, from hydrolysing complex carbohydrates to primary and secondary fermenters that produce short-chain fatty acids and species that use hydrogen for their metabolism [39]. However, simple sugars like glucose are universally possible to assimilate and energetically cheap to metabolise, which makes them an evolutionarily preferred carbon source when available [40]. When such substrates are abundant, most community members can independently access energy without relying on complementary metabolic pathways provided by other microbes. This leads to a breakdown in mutualistic and commensal interactions, as cross-feeding and metabolic interdependence become less advantageous. As a result, community dynamics shift toward individualism, with species competing more directly for the same resources, thereby fostering exploitative interactions and reducing ecological cohesion. A consistent driver across these interaction shifts is their association with nutrient intake, particularly simple sugars. In the FoCus cohort, consumption of glucose and fructose is associated with an increase in exploitative interactions, suggesting that these rapidly metabolised substrates may amplify microbial competition or antagonism for shared resources. In contrast, consumption of vitamins, nitrogen-containing compounds, polyamines, and tryptophan promoted collaborative interactions, particularly commensalism. This is in line with the notion that those compounds are expensive to build up or must be consumed by the environment as public goods [39].

Together, our findings highlight a consistent ecological signal across disease, lifestyle, and diet, which is why dietary modulation may offer a strategy for restoring cooperative microbial interactions in dysbiotic or industrialised microbiomes. However, this potential remains constrained by modelling limitations.

The intervention analysis highlights that targeted nutrient supplementation can modulate the ecological interaction structure of the gut microbiome, but these effects are not always aligned with observational associations from dietary intake. For example, while higher habitual intake of simple sugars (glucose and fructose) was linked to an increase in exploitative interactions in cohort analyses, supplementation of these same sugars in silico increased collaborative interactions. This apparent contradiction likely reflects fundamental differences between dietary patterns and supplementation scenarios in a controlled metabolic model. In the real world, sustained high intake of simple sugars may reduce resource diversity and increase competition for easily accessible substrates, promoting exploitative dynamics. By contrast, supplementation in a model context can create an abundance of a readily metabolizable carbon source for both species, reducing competitive pressure. This “dose” dependency in turn increases their predicted growth rates and would be later interpreted as an increase in collaboration.

Vitamins and purines consistently promoted collaborative dynamics in silico, supporting the view that micronutrients and nucleotides act as public goods within microbial communities [41, 42]. These compounds often require specialised biosynthetic pathways, so supplementation may relieve metabolic bottlenecks and encourage positive-sum exchanges, reinforcing cooperative interactions. Besides, we observe that nitrogen-containing compounds (cytidine, deoxyadenosine, guanosine, xanthine, urate, deoxyguanosine, hypoxanthine, and adenosine) are boosting collaborative interactions. This is particularly interesting if we consider that the colon, except towards its end, is usually a nitrogen-depleted environment [39]

Collectively, these findings emphasise that nutrient availability strongly governs microbial interaction patterns, but the effect direction can depend on abundance context, metabolic accessibility, and whether resources are universally utilisable or pathway-specific. This suggests that dietary interventions aimed at restoring microbial cooperation must consider not only nutrient type but also systemic context and metabolic redundancy within the community.

Currently, EcoGS models only pairwise interactions, which may fail to capture emergent behaviours in more complex microbial consortia. Although higher-order modelling is theoretically possible, the combinatorial explosion, e.g., ∼89 million triplet interactions for all strains in the AGORA gut collection, renders it computationally infeasible with current approaches. Moreover, precision diet modelling is limited by the need to re-run simulations for each specific diet, which is computationally intensive. Finally, the reliance on gap-filling algorithms during genome-scale reconstruction, via gapseq, may reduce the number of unique metabolic exchanges, biasing interaction classifications toward competition. In summary, EcoGS represents a valuable tool for investigating microbial ecology at scale. Its ability to uncover consistent interaction shifts across human populations and dietary patterns underscores the microbiome’s role as a mediator and driver of host-environment interactions. Future developments, such as scalable multi-species modelling, dynamic nutrient flux integration, and tighter coupling with empirical validation, will further enhance its utility in both ecological theory and microbiome-based intervention design.

## 7. List of abbreviations

EcoGS: The presented tool to predict the **Eco**logical interactions based on **G**enome-**s**cale metabolic models
GMbC: The **G**lobal **M**icro**b**iome **C**onservancy
EIR: In this document refers to the **E**cological **I**nteraction **R**atios
FDR: **F**alse **D**iscovery **R**ate, a method for correcting for multiple testing
BMI: **B**ody **M**ass **I**ndex, calculated by dividing the weight by the height squared

## 8. Declarations

### 8.1. Ethics approval and consent to participate

Permission to utilise the FoCus cohort data, which included ethics approval, was obtained by submitting a structured online application (available at https://www.uksh.de/P2N/) to the Kiel 2.0 Network (Reference Number: 2024-009). Permission to utilise the GMbC cohort data was obtained by the co-authors, GMbC consortium leaders, and science & ethics advisory board members Mathieu Groussin and Mathilde Poyet. Research & ethics approvals were obtained from the MIT IRB (protocol #1612797956) and from the Ethics Commission of the Medical Faculty of Kiel University (Studie D 511/24). Permits were also obtained in each sampled country prior to the start of sample collection, from the following ethics committees:

● Cameroon: Comité National d’Ethique de la Recherche pour la Santé Humaine, protocol #2017/05/901/CE/CNERSH/SP;
● Canada: Nunavut Research Institute, protocol #0205217N-M;
● Central African Republic: Comité d’Éthique et Scientifique de la Faculté des Sciences de la Santé de l’Université de Bangui, No. 17/UB/FACSS/CES20;
● Finland: Coordinating Ethics Committee of Helsinki and Uusimaa Hospital District, protocol #1527/2017;
● Ghana: Cape Coast Teaching Hospital Ethical Review Committee, protocol #CCTHERC/RS/EC/2016/3 and Committee on Human Research, Publication and Ethics of the Komfo Anokye Teaching Hospital, protocol #CHRPE/AP/398/18;
● Malaysia: Universiti Malaya Medical Research Ethics Committee, MREC ID No.: 2018219-6033;
● Nigeria: National Health Research Ethics Committee of Nigeria, protocol #NHREC/01/01/2007-29/04/2018.
● Rwanda: National Ethics Committee, protocol IRB 00001497 of IORG0001100
● Senegal: Comité National d’Ethique pour la recherche en Santé, No.: 0000073 MSAS/DPRS/CNERS
● Tanzania: National Institute for Medical Research, protocol #NIMR/HQ/R.8a/Vol. IX/2657;
● Thailand: The Human Research Ethics Committee of the Faculty of Medicine of Thammasat University No.1, Certificate of Approval 143/2019
● United States of America: Chief Dull Knife College, protocol #FWA00020985;

### 8.2. Availability of data and materials

The statistical analysis and plotting R scripts can be found under: https://doi.org/10.6084/m9.figshare.30149005

## 8.3. Acknowledgements

We thank the following funding organisations

‒ Excellence Cluster for Precision Medicine in Chronic Inflammation (PMI)
‒ The German Research Foundation within the collaborative research centre “Origin and Function of Metaorganisms,” CRC1182
‒ Research Group miTarget
‒ iTREAT
‒ DFG project ExoMod

We thank Dr Johannes Zimmermann and Dr Robin Koch for the discussions and support

## 9. Supplementary material

**Figure S1:**
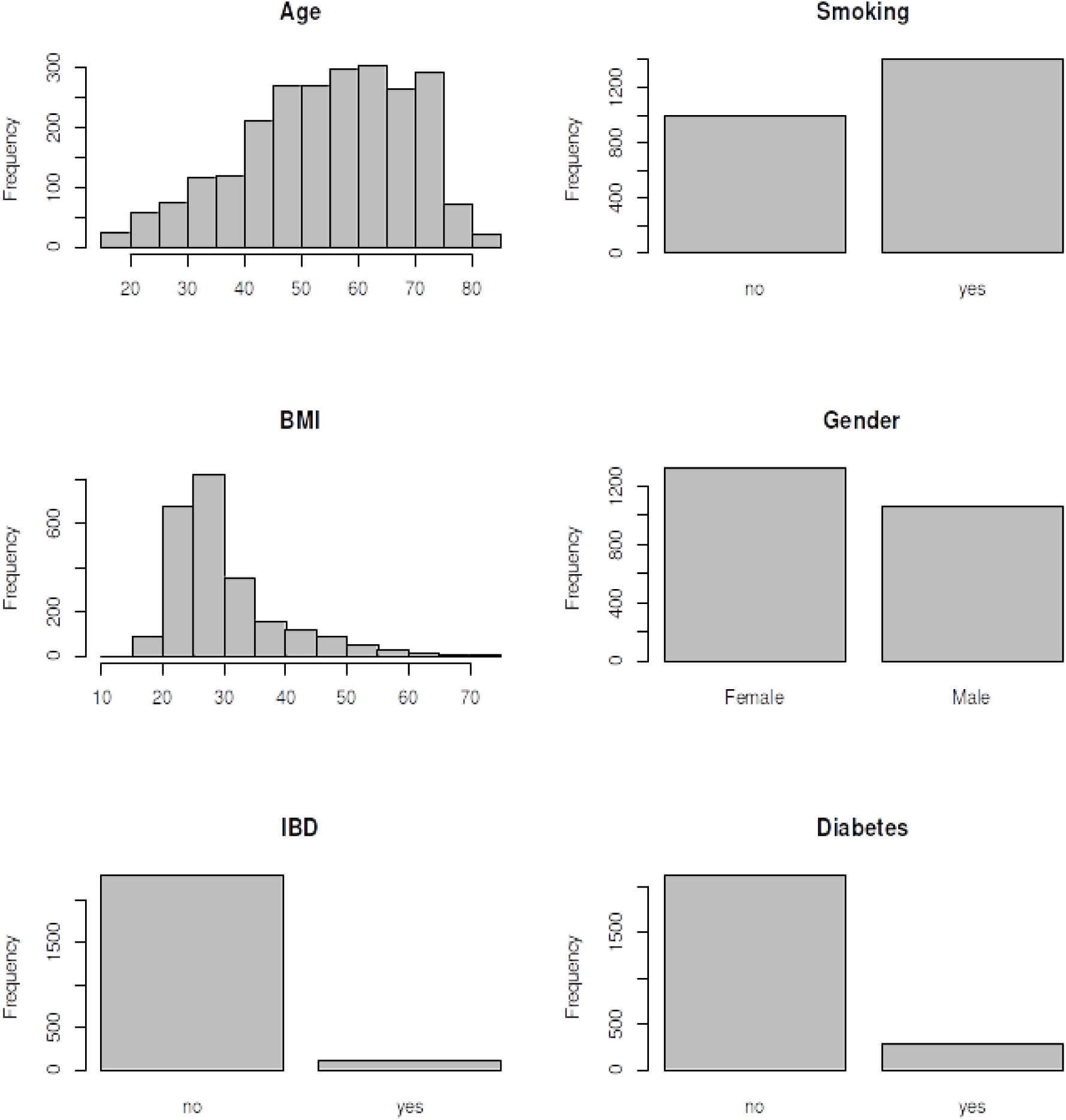
The distribution of FoCus cohort descriptive phenotypic data

**Figure S2:**
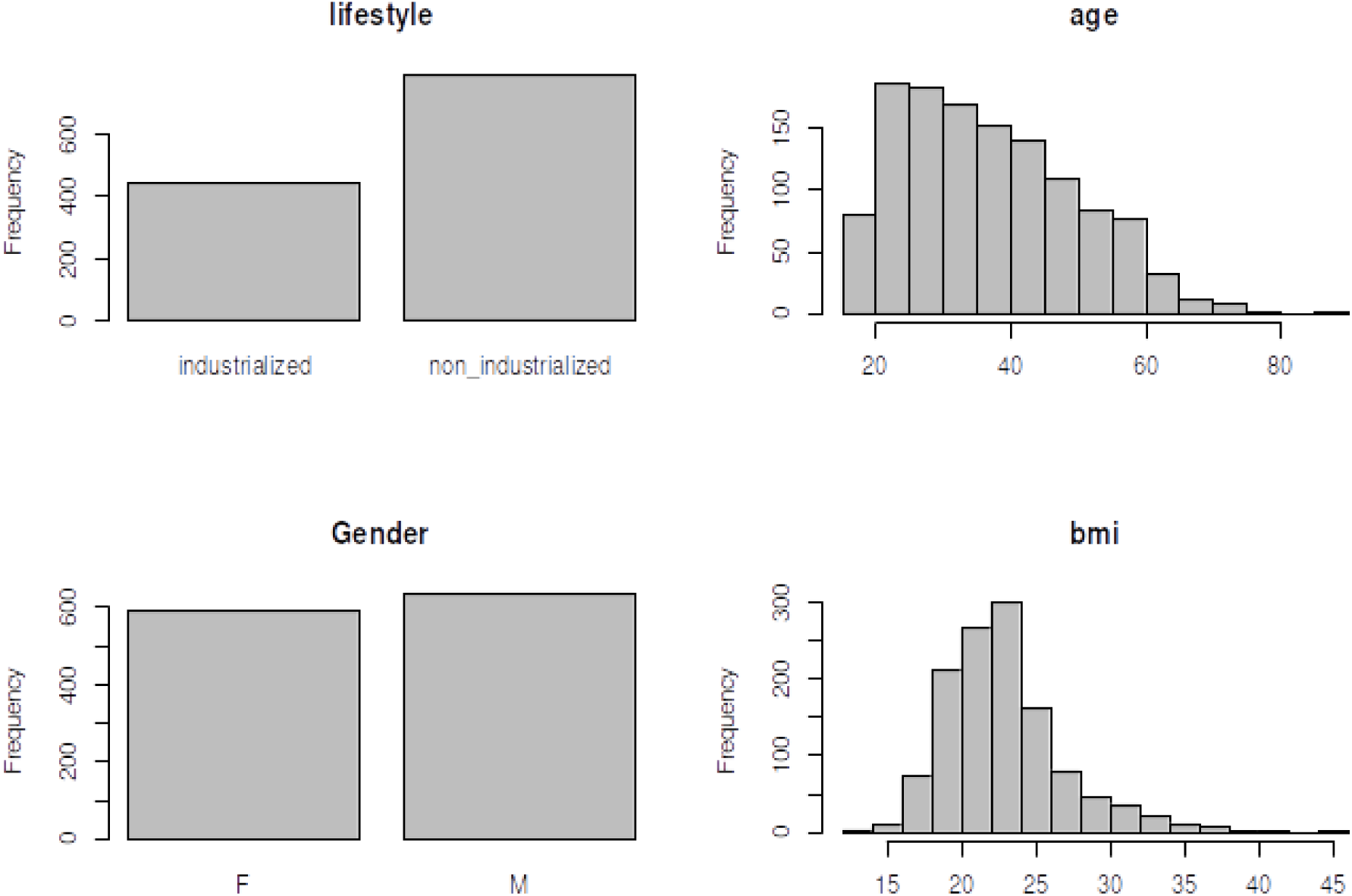
The distribution of GmBC cohort descriptive phenotypic data

Supplementary table S1:

https://docs.google.com/spreadsheets/d/1IXCp2RYRWFko31_K3P7fXAb4CcNPRfxyJunhAKgZyx4/edit?usp=sharing

## Notes

https://figshare.com/articles/software/R_scripts_for_statistics_and_plotting_that_were_used_to_generate_the_Pvalues_and_plots_published_in_the_EcoGS_manuscript/30149005

http://www.github.com/KaletaLab/EcoGS

### Competing Interest Statement

The authors have declared no competing interest.

